# Decoupling nucleolar stress from DNA-damage in HCT116 colon cancer cells by targeting the interaction between ribosomal assembly factors Bop1/WDR12

**DOI:** 10.64898/2026.03.11.711022

**Authors:** Susana Masiá, Jerónimo Bravo

**Affiliations:** Instituto de Biomedicina de Valencia, Consejo Superior de Investigaciones Científicas (CSIC), c/ Jaime Roig 11, 46010 Valencia, Spain

## Abstract

Ribosome biogenesis is a hallmark of cancer and has emerged as an effective target for treatment. This study investigates novel strategies in cancer therapy by disrupting specific protein-protein interactions essential for the ribosome biogenesis process. Using peptide mimetics derived from human Bop1, conjugated with cell-penetrating sequences, we assessed their binding affinity, cellular internalization, and biological effects in HCT116 colon cancer cells. The peptides disrupted Bop1-WDR12 interaction, leading to nucleolar stress, reduced ribosome biogenesis and decreased protein synthesis. Importantly, these interventions triggered apoptosis via caspase activation without eliciting DNA double-strand breaks. These findings demonstrate that targeted disruption of ribosome assembly pathways can induce cancer cell death independently of genotoxicity, offering a promising non-genotoxic therapeutic approach that would minimize side effects. Targeting protein-protein interactions of assembly factors during ribosome maturation can be a new strategy with significant potential to improve cancer treatments by selectively impairing tumor growth while not contributing to genomic instability.

## INTRODUCTION

Ribosomes are essential molecular machines that synthesize proteins. The mature 80S eukaryotic ribosome is formed by four ribosomal RNAs and 80 ribosomal proteins organized as the 40S or small subunit and the 60S or large subunit. The process of assembling the ribosome is known as the ribosome biogenesis process and is initiated at the nucleolus with the transcription of rRNA by RNA pol I ^1,2^. The biosynthesis of ribosomes is an essential, evolutionary highly conserved and energetically demanding process involving the coordinated assembly of numerous proteins and RNA molecules. A large number of assembly factors, which will be absent from the functional ribosome, are required at different stages during the maturation process for the splicing and processing of the pre rRNA including RNA processing factors, protein chaperones, GTPases or RNA helicases. Modifications in the complex multistep mechanism of ribosome biogenesis precipitate nucleolar stress ^3^.

Over the past decade, the field has evolved from recognizing ribosome biogenesis as a hallmark of cancer to its identification as a promising target for novel therapies ^4–7^ mostly due to its fundamental role in supporting increased protein production, uncontrolled cell proliferation and tumor progression ^6,8^. Rapid division and metabolic adaptation exhibited by cancer cells demand high protein production, making ribosome biogenesis a key regulatory process that is clearly increased compared to normal cells ^9^. RNA pol I is under the control of both tumor suppressor genes (including p53, Rb and Arf) and oncogenes (including MYC, MAPK/ERK, PI3K and AKT). Myc and PTEN act as master regulators of ribosome biogenesis, translation control and protein synthesis as a result ^10^, while tumor suppressors like p53 promote its suppression. Another indication of its therapeutic potential is that cell proliferation could be blocked by inhibiting the production of new ribosomes ^4,11^. In fact, it has been proposed that tumor cells are addicted to increased ribosome production since inhibition is generally cytotoxic in proliferating cells ^5^ indicating that cancer cells, somehow, become dependent on ribosome synthesis. Disruption of ribosome biogenesis can lead to the inhibition of protein synthesis, cell growth and proliferation and has therefore received much attention as an emerging potential therapeutic target for cancer treatment ^12^. Interestingly it has also been shown that ribosome biogenesis inhibition reduces metastatic seeding ^13^. An additional rationale for evaluating this mechanism as a potential therapeutic target resides in the acknowledgment that the molecular interactions of various alkylating, intercalating molecules, and kinase inhibitors, which are presently employed in oncological therapies, contribute to their efficacy against neoplastic cells through the suppression of ribosome synthesis. Agents such as oxaliplatin, doxorubicin, and methotrexate have been demonstrated to impede rRNA transcription at concentrations that compromise nucleolar integrity, whereas compounds like camptothecin, flavopiridol, and roscovitine inhibit preliminary rRNA processing events. Concurrently, 5-fluorouracil (5-FU) and homoharringtonine engage later stages, resulting in nucleolar distress and the subsequent activation of the compromised ribosome biogenesis checkpoint ^14,15^. These pharmacological agents, originally developed to induce DNA damage or obstruct various cellular mechanisms, are presently acknowledged for their role in enhancing antitumor efficacy through the inhibition of ribosome synthesis ^16,17^. Nevertheless, mechanistic investigations are revealing that a majority of these therapeutic molecules demonstrate dose-dependent pleiotropic effects and exhibit a significant degree of non-specificity. A more deliberate and targeted strategy is the recent exploration of RNA polymerase I inhibitors, which has resulted in the identification of several novel compounds with noteworthy anti-cancer properties, such as CX-5461, which is currently undergoing phase I/II clinical trials ^18^. In a manner analogous to the current chemotherapeutic agents targeting ribosome biogenesis, the inhibition of RNA polymerase I specifically addresses the preliminary phase of ribosomal RNA synthesis and has also been correlated with the occurrence of DNA double-strand breaks ^19,20^. Consequently, notwithstanding its status as a dynamic and promising avenue of inquiry, the pursuit of innovative or enhanced inhibitors of ribosome biogenesis should persist.

Recent progress in the field of structural biology has elucidated molecular intricacies associated with various phases of ribosome biogenesis pathways in both yeast and human ^21,22^. This acquired knowledge may enable the formulation of more precise therapeutic interventions, potentially permitting the suppression of ribosome production while avoiding the induction of DNA damage. The so called PeBoW complex is formed by Pes1, Bop1 and WDR12 (Nop7, Erb1 and Ytm1 respectively in yeast) and participates in the maturation of the ribosomal 60S ribosomal subunit. They belong to the A_3_ family of assembly factors and are involved in the removal of the Internal transcribed Spacer 1 between the 18S and 5.8S rRNA in the precursor pre-rRNA. Bop1 interacts directly with WDR12 and have both been implicated in cancer progression ^23–25^. WDR12 has emerged as a promising candidate for both cancer diagnosis and therapeutic targeting due to its significant role in various malignancies. Dysregulation of WDR12 expression has been observed in multiple cancer types, including hepatocellular carcinoma ^26^, glioblastoma ^27,28^ and colorectal cancers ^29^. Its expression levels correlate with patient prognosis, suggesting its potential as a biomarker ^25,28^. Functional studies employing lentivirus-mediated knockdown of WDR12 in CRC cell lines such as HCT116 and SW620 show a marked decrease in cell proliferation and viability in parallel with an increase in apoptosis ^29^.

We have previously articulated the structural properties of the Erb1-Ytm1 complex, which corresponds to the orthologous counterparts of human Bop1-WDR12, respectively. We illuminated the molecular complexities inherent to the interacting domains and demonstrated that modifications to this interface result in the impairment of ribosome biogenesis and cellular proliferation in Saccharomyces cerevisiae ^30^. The examination of the interacting interface of the Chaetomium thermophilus Erb1 C-terminal domain in conjunction with Ytm1 facilitated the development of peptides intended to disrupt the formation of the complex ^31^. Recent Cryo-Electron Microscopy structures of pre-60S intermediates elucidated the significance of the Bop1-WDR12 interaction in the excision of internal transcribed spacers from the pre-rRNA molecule, as well as the considerable degree of structural and evolutionary conservation observed ^22^.

In this study, we delineate peptides that specifically target the Bop1-WDR12 interaction, demonstrating their capacity to diminish cellular viability, attenuate protein synthesis, and promote apoptosis independently of genotoxicity in HCT116 colon cancer cells.

## RESULTS

### Biotinilated BOP1 derived peptides bind WDR12

In a prior investigation, we evaluated a preliminary collection of six peptides originating from the interface between Erb1 and Ytm1 ^31^. Two peptides that were extracted from the Erb1 sequence exhibited superior performance in both competitive and biophysical binding assays. Considering the substantial level of evolutionary conservation, we evaluated the human Bop1 variants of these two peptides for its binding affinity with WDR12. Biot-Bop1_439-449_ and Biot-Bop1_725-734_ feature a biotin moiety strategically positioned at the N-terminus of peptides that encapsulate respectively human Bop1 residues 439-449 (LLWEVATARCVR, which encompasses a segment of strands 1c to 2c of the β-propeller) and 725-734 (IFHPTQPWVF), which encapsulates part of strands 7a and 7b and the loop between them).

Parameters for Biot-Bop1_405-412_ couldn’t be obtained due to solubility problems of the peptide at higher concentrations. Biot-Bop1_439-449_showed a Kd in the low micromolar range, similar to the affinities obtained when using the *Chaetomium thermophilum* Erb1 derived peptides and Ytm1 ^31^ (Fig1A).

### ex vivo effect of Bop1 derived peptides

#### Cell internalisation of WDR12 binding peptides

We aimed to assess the impact of the peptides in HCT116 colon carcinoma cells. To facilitate the internalization of the selected sequences into the cellular environment, the peptide GRKKRRQRRRPQ, which is derived from the transactivator of transcription (TAT) of the human immunodeficiency virus, was conjugated to the N-terminus of Bop1 peptides 439-449 and 725-734 generating P10hs and P11hs peptides respectively. N-terminal FITC-conjugated P10hs and P11hs peptides were evaluated for their ability to penetrate cellular membranes and internalize within the cellular structure, leading to the discovery of a remarkably extensive distribution throughout the entirety of the cellular architecture, which notably includes the nucleolus as graphically depicted in Figure 1B.

**Figure 1.**
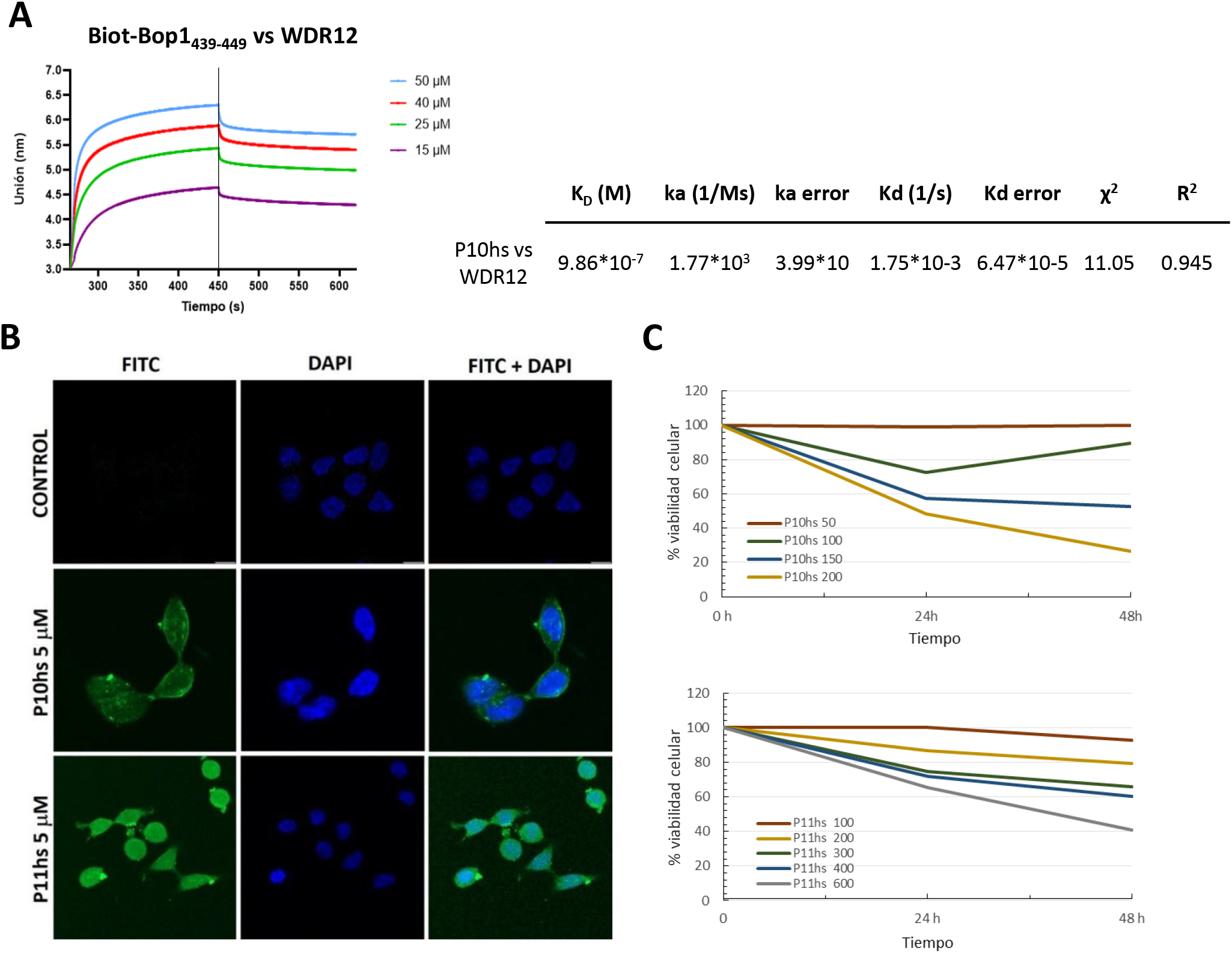
Bop1_439-449_ in vitro binding with WDR12, FITC-P10hs and FITC-P11hs cellular localization and HCT116 viability in the presence of P10hs and P11hs. **(A)** Biolayer Interferometry assay. Streptavidin sensor tips loaded with Biot-Bop1_439-449_ using WDR12 as analyte. **(B)** Confocal microscopy images of HCT116 cells treated for 24 h with 5 µM FITC-labeled P10hs and P11hs peptides (green). DAPI was used as a nuclear marker (blue). Scale: 10 µm. **(C)** Quantification of HCT116 cell viability after treatment with 50, 100, 150, and 200 µM of P10hs (upper panel) and 100, 200, 300, 400, and 600 µM of P11hs at 24 and 48 hours (lower panel).

#### Penetrating P10hs and P11hs decrease HTC116 cell survival rates

Following this initial assessment, we proceeded to conduct a thorough investigation into the survival rates of the cells across a variety of temporal intervals and peptide concentrations, specifically at time points of 0, 24, and 48 hours, as illustrated in Figure 1C. The analysis of survival rates demonstrated a significant decline that occurred in a manner that was dependent on the concentration of the peptides for both P10hs and P11hs when compared to the baseline control provided by the standard TAT sequence. Notably, it was observed that P10hs requires a lower concentration to achieve the half-maximal inhibitory concentration (IC50), thereby indicating that it possesses superior efficacy over P11hs in diminishing the viability of HCT116 cells (Fig 1C).

#### P10hs and P11hs disrupt nucleolar integrity and produce overall protein synthesis reduction in HTC116 cells

The unusual and aberrant localization of the nucleolar protein known as nucleolin, which is a crucial factor in the nucleolus, is intricately correlated with significant structural disruptions occurring within the nucleolus itself, along with a marked reduction in the functionality of RNA polymerase I (Pol I), and an impairment in the maturation processes of ribosomal RNA (rRNA). Following an exposure lasting 24 hours to the specified treatments P10hs and P11hs, nucleolin demonstrates a remarkably widespread distribution throughout the nucleoplasm (Fig 2), a state that sharply contrasts with the untreated or penetrating sequence TAT treated HCT116 cells, which display a typical and anticipated nucleolar localization, thus indicating nucleolar stress and a profound disruption in the fundamental and crucial processes that regulate ribosome biogenesis.

**Figure 2.**
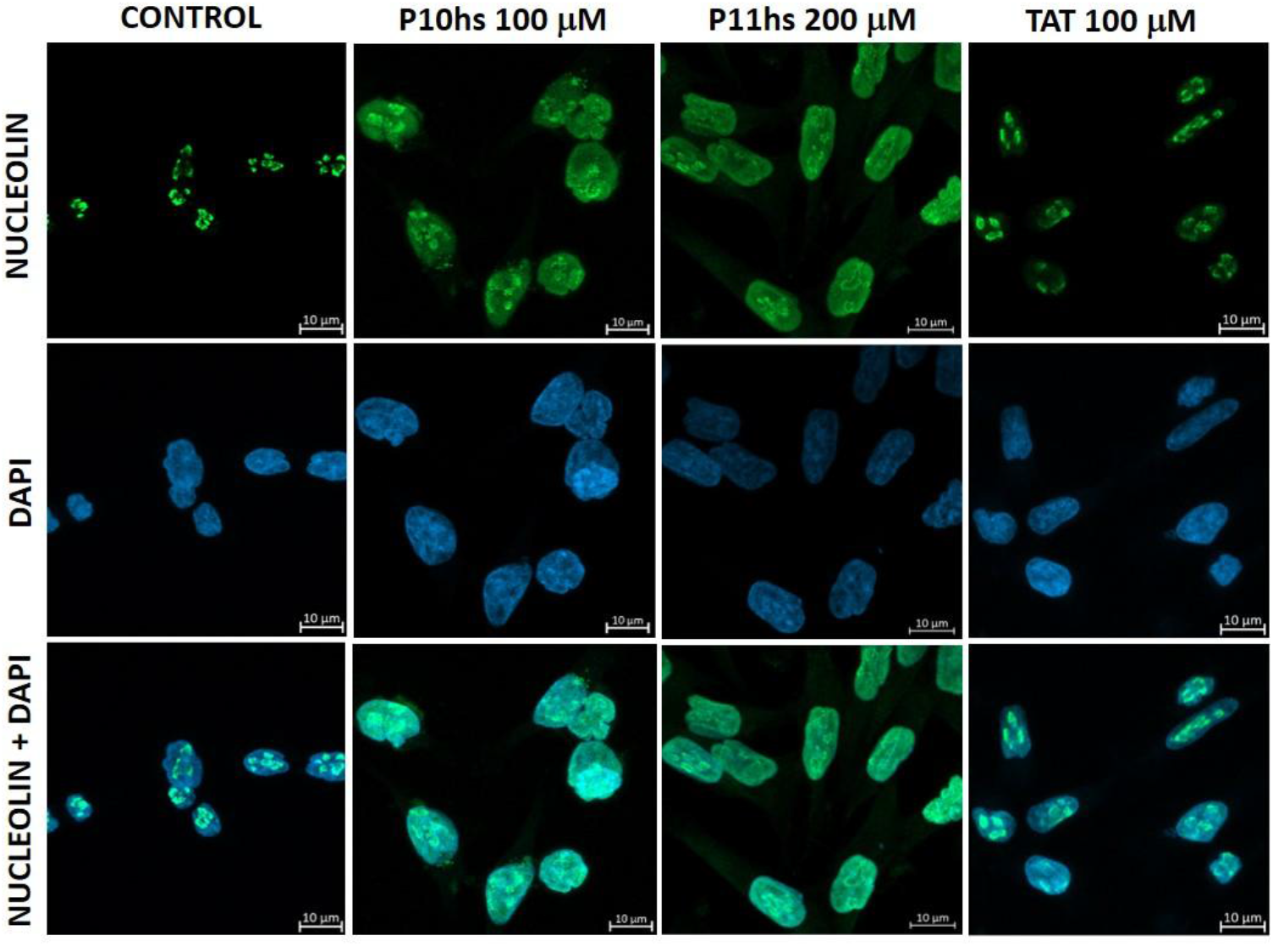
Delocalization of nucleolin after treatment with P10hs and P11hs peptides. Confocal microscopy images of untreated HCT116 cells (CONTROL) and cells treated with 100 µM P10hs, 200 µM P11hs, or 100 µM TAT for 24 h. Nucleolin is clearly localized in granular nucleosomes in both untreated cells and cells treated with the TAT peptide (green). In contrast, it is spread throughout the nucleus in cells treated with P10hs and P11hs. Nuclei were stained with DAPI (blue). Scale: 10 µm.

Moreover, the polysome profiles derived from P10hs and P11hs HCT116 treated cells, exhibit a significant reduction in the 80S fraction, as illustrated in Fig 3A, thereby underscoring the adverse effects of the peptide treatments on ribosomal assembly and functionality. This specific finding suggests possible ramifications for translational processes following P10hs or P11hs administration. In fact, a 24-hour exposure of HCT116 cells to either P10hs or P11hs reveals detrimental implications for the process of protein biosynthesis (Fig 3B). A quantification of puromycin-tagged polypeptides conducted using the SUnSET methodology illustrates a significant decline of more than 40% in overall protein synthesis (Fig 3C), thereby establishing a correlation between the previously observed decrease in cellular viability (Fig 1C) and the reduction in protein biosynthesis.

**Figure 3.**
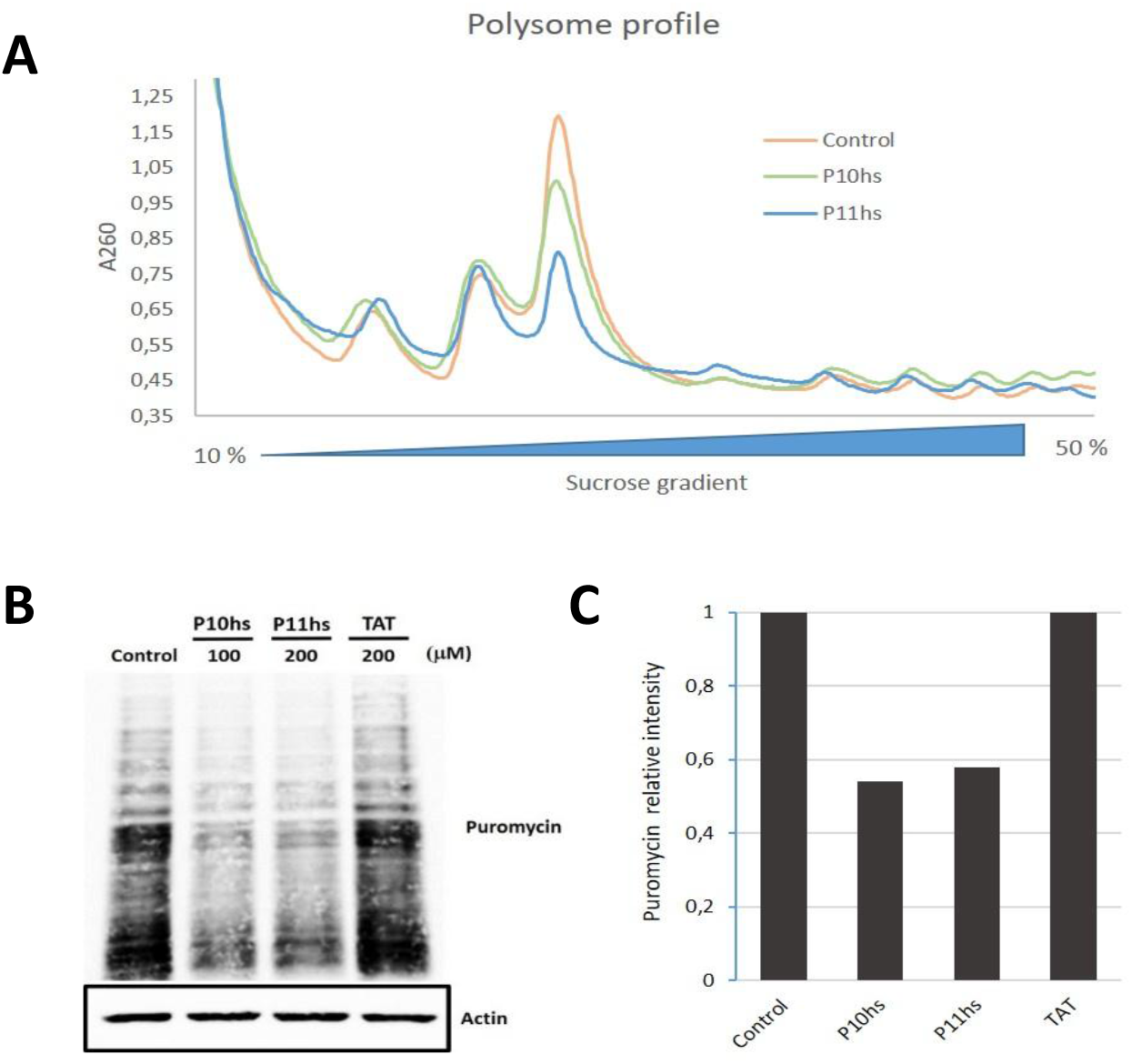
Reduction in protein synthesis following treatment with peptides P10hs and P11hs. **(A)** Polysomal profiles derived from HCT116 cells treated with P10hs and P11hs showing a significant reduction in the 80S fraction. HCT116 cells were treated with 100 µM P10hs or 400 µM P11hs for 24 hours. Untreated cells were used as a control. The corresponding cytosolic extracts were sedimented by centrifugation in 10-50% sucrose gradients. **(B)** SUnSET assay in which HCT116 cells were treated or not treated for 24 hours with 100 µM of the P10hs peptide, 200 µM of the P11hs peptide, or 200 µM of the TAT peptide and exposed to puromycin (5 μg/mL) during 15 minutes prior to protein extraction. Protein translation rates were assessed by Western blot analysis using an anti-puromycin antibody. Actin was used as a loading control. **(C)** Quantification of the SUnSET assay result using ImageJ software.

#### P10hs and P11hs *promote apoptosis in HTC116 cells*

Upon demonstrating the internalization of the cell-penetrating variants of the P10hs and P11hs peptides, their effect on cell viability alongside their involvement in the disruption of ribosome biogenesis and protein synthesis, our intention is to further elucidate the cellular ramifications of neoplastic cells subjected to treatment with these peptides.

Following our demonstration of the internalization of cell-penetrating variants of both Bop1-derived peptides, as well as their role in disrupting ribosome biogenesis and protein synthesis, we aim to examine deeper into the cellular consequences experienced by neoplastic cells treated with these peptides. To achieve this, we employed flow cytometry, utilizing Annexin V-FITC and propidium iodide (PI) as standard reagents for quantifying the various stages of cell death that offers a robust method for distinguishing between live, early apoptotic, late apoptotic, and necrotic cells, thereby providing a comprehensive overview of cellular viability and death pathways. In our experiments, we administered the peptides to HCT116 cells and observed significant differences in late apoptosis and necrosis at the 24-hour mark, when comparing the effects of 100 or 150 μM P10hs and 200 or 400 μM P11hs against control groups (Fig 4A, quadrant Q2). A dose effect in the apoptotic events is observed for both peptides. In accordance with the cellular viability results (Fig 1), P11hs necessitates a heightened concentration in comparison to P10hs to elicit apoptotic responses in HTC116 cells. Notably, the treatment of HCT116 cells with TAT at a concentration of 400 μM resulted in cellular outcomes that were comparable to those of untreated cells (Fig 4A top). This finding implies that the observed differences in peptides P10hs and P11hs treated cells with respect untreated HTC116 cells cannot be attributed to the cell-penetrating sequences shared by both peptides.

**Figure 4.**
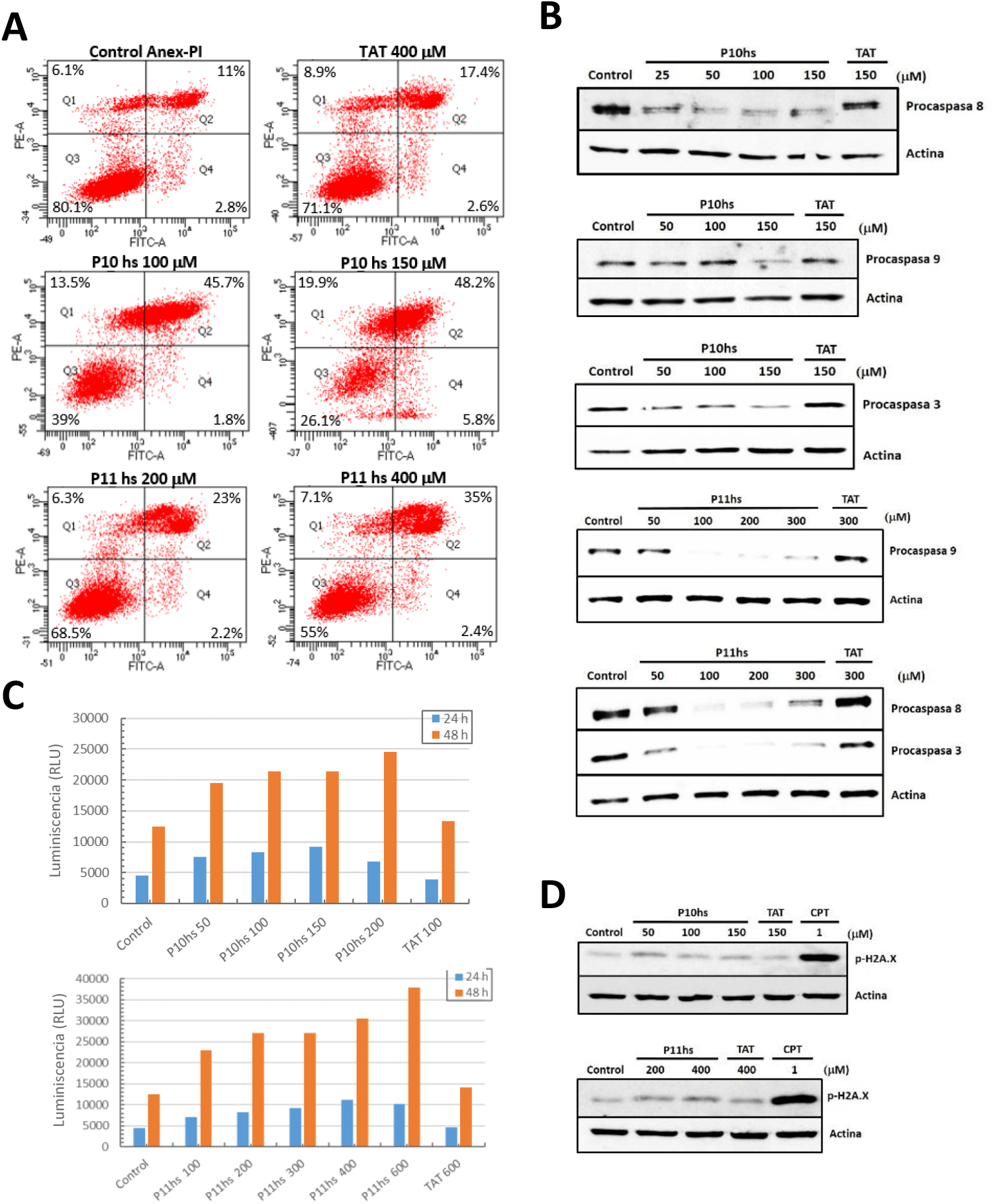
Apoptosis activation in HCT116 cells after treatment with P10hs and P11hs peptides. **(A)** Detection of different cell populations after 24 h treatment with penetrating peptides using Annexin FITC V/PI staining by flow cytometry. Representative graphs of double staining with Annexin-FITC V/PI of untreated HCT116 cells and after treatment with 100 and 150 μM P10hs; 200 and 400 μM P11hs; and 400 μM TAT. Each quadrant shows the mean percentage of live cells (Q3: Annexin FITC V-/PI-); early apoptotic cells (Q4: Annexin FITC V+/PI-); late apoptotic cells (Q2: Annexin FITC V+/PI+); and necrotic cells (Q1: Annexin FITC V-/PI+). **(B)** Caspase activation after treatment with P10hs and P11hs peptides in HCT116 cells. Cells were treated with different concentrations of P10hs (25, 50, 100, and 150 µM), P11hs (50, 100, 200, and 300 µM), and TAT (150 and 300 µM) for 24 hours. Total lysates from HCT116 cells were analyzed by Western blot using specific antibodies against the proactive forms of caspase 3, caspase 8, and caspase 9, with actin as a loading control. **(C)** Quantification of caspase 3/7 activity in HCT116 cells relative to the number of viable cells after treatment with P10hs (50, 100, 150, and 200 µM), P11hs (100, 200, 300, 400, and 600 µM), and TAT (100 and 600 µM). All treatments were performed for 24 and 48 hours. RLU: relative light units. **(D)** DNA integrity after treatment with the peptides P10hs, P11hs, and TAT. Lysates from HCT116 cells were analyzed by Western blot using the γ-H2A.X antibody as a marker of DNA damage and actin as a loading control. CPT: camptothecin.

In order to enhance our comprehension of the cellular death mechanisms in HCT116 cells triggered by P10hs and P11hs treatment, we conducted a western blot analysis utilizing antibodies specific to procaspases alongside caspase activation luminescence assays. As depicted in figure 4B, we observe a statistically significant reduction in procaspases 3, 8 and 9, which indicates caspase activation following a 24-hour exposure of HCT116 cells to P10hs or P11hs (Fig 4B). This is further confirmed by activation of the proluminescent caspase-3/7 DEVD-aminoluciferin cleavage in a luminescent assay. HTC116 P10hs and P11hs 24h or 48h treatment results indicate that caspase 3/7 activation is detected in as little as 50 μM and 100 μM respectively (Figure 4C). The findings of our investigation suggest that the late stages of apoptosis observed in the cell cytometry assays are associated with the activation of executioner caspases, specifically caspase 3, in conjunction with initiator caspases such as caspase 8 or caspase 9, subsequent to the administration of 24h and 48h treatments P10hs and P11hs on HTC116 cells.

#### HCT116 apoptosis induced by P10hs and P11hs is not mediated by DNA damage

New compounds known to influence ribosome biogenesis, such as CX-5461, along with traditional chemotherapeutic agents like doxorubicin, induce DNA damage that results in adverse side effects ^32^. Consequently, we sought to examine whether the cytotoxic effects exhibited by P10hs and P11hs were mediated through DNA damage. The phosphorylation of the histone variant H2AX at the Ser139 residue to produce γ-H2AX is extensively employed for the detection and quantification of DNA damage and genomic instability, as it exhibits a strong correlation with double-strand DNA breaks.

Camptothecin (CPT), are recognized for its ability to induce double-strand breaks that elicit cell cycle arrest and subsequent apoptosis, and have been employed as a positive control in experimental assays. Western blot analyses utilizing antibodies specific to γ-H2AX reveal comparable levels for the 24-hour treatments of P10hs and P11hs as those observed in untreated cells, which serve as a negative control (FIG 4D upper panel). Treatment of HTC116 cells with 50, 100, and 150 μM concentrations of P10hs for a duration of 24 hours does not elicit a statistically significant increase in γ-H2AX levels. Likewise, HTC116 cells subjected to 24-hour treatments with 200 and 400 μM concentrations of P11hs exhibited γ-H2AX levels that were comparable to those of untreated cells (FIG 4D lower panel). Similarly to P10hs and P11hs, the penetrating sequence TAT at 150 or 400 μM didn’t show an increase of γ-H2AX. These findings suggest that the apoptosis observed in HTC116 cells following treatment with P10hs and P11hs is not attributable to double-strand DNA damage, in contrast to other agents reported to target ribosome biogenesis.

## DISCUSSION

In a similar manner to our prior research utilizing Chaetomium thermophilum Erb1_Ytm1 sequences ^31^, we have now demonstrated that P10hs originating from human BOP1 exhibit *in vitro* interaction with WDR12 within a low μM concentration range. The cell-penetrating peptidic variants P10hs and P11hs possess the capacity to infiltrate HTC116 cells, inducing nucleolar stress and causing alterations in ribosome profiling, analogous to the interface modifications executed using yeast ^30^ thereby underscoring the significant degree of structural and functional conservation that has transpired throughout evolution in the context of eukaryotic ribosome biogenesis. This also emphasizes the utility of yeast as a model organism for elucidating the mechanisms underpinning ribosome synthesis, a fact that has been previously recognized.

Interference peptides designed to disrupt the BOP1/WDR12 interaction induce a variety of cellular ramifications, including diminished cell viability, the onset of nucleolar stress, and a decrease in protein biosynthesis. This observation aligns with the downregulation of ribosome biogenesis, as nucleolar stress represents a pathological state resulting from disturbances within this pathway, ultimately culminating in cell cycle arrest or programmed cell death. The observed decrease in HTC116 cell viability following the administration of P10hs and P11hs corresponds to an apoptosis triggering, as evidenced by the concomitant activation of initiator caspases 8 and 9, and effector caspase 3/7, which serves as a definitive indication of the engagement of apoptotic pathways. This is also in agreement with the well-established relationship between ribosome biogenesis impairment and apoptotic events. The induction of caspases 8 and 9 subsequent to the application of P10hs and P11hs on HCT116 cells may imply the participation of both the extrinsic and intrinsic apoptotic pathways at a defined phase; nevertheless, additional research is necessary to comprehensively dissect this hypothesis.

It has been reported that double strand breaks (DSBs) and genotoxic stress may arise in conjunction with the inhibition of ribosome biogenesis, particularly in the context of conventional chemotherapeutic agents reported to mechanistically affect ribosomal synthesis. Several current therapeutic agents—such as actinomycin D, cisplatin, oxaliplatin, 5-fluorouracil, doxorubicin, mitoxantrone, camptothecin, or etoposide simultaneously target ribosome biogenesis and produce DNA damage. Other agents currently under development like CX-3543 and CX-5461 show, to this regard, a similar behaviour. It can be discussed whether this dual action contributes to their anticancer efficacy by engaging both nucleolar stress and DNA damage response pathways. However, genotoxicity induces mutations that contribute to tumor recurrence and secondary malignancies, further exacerbating patient morbidity, especially in pediatric populations^33 34^. Inflicting DNA damage to induce cell death leads to adverse effects such as gastrointestinal dysfunction, alopecia, and premature aging^35^. There exists, consequently, a real need for novel therapeutics that minimize or uncouple DNA damage from nucleolar stress, thereby reducing the likelihood that double strand breaks formation will compromise their clinical utility. During the progression of our research endeavors, the nucleolar stress elicited by P10hs and P11hs in HTC116 cellular models instigates apoptosis while concurrently failing to promote the occurrence of DNA double strand breaks, thereby paving the way for an innovative era of non-genotoxic anticancer therapeutic strategies.

### Ribosome biogenesis impairment without targeting rDNA transcription

In this context, it is imperative to underscore that nearly all endeavors aimed at the attenuation of ribosomal synthesis effectively accomplish their objectives by focusing on the initial phases of the process through the modification of rDNA or the interruption of rRNA transcription. Conversely, instead of predominantly targeting either pathway of rRNA synthesis, we present evidence that intervention in subsequent stages of ribosome biogenesis is feasible for the purpose of disrupting ribosome biogenesis. Moreover, this promotion of nucleolar stress has been realized without the induction of DNA damage, thereby offering novel avenues for a targeted approach to cancer treatment. It is therefore our assertion that instead of focusing exclusively on the transcriptional processes of ribosomal RNA (rRNA), the deliberate targeting of subsequent protein translation events taking place during the ribosome biogenesis pathway, facilitates a disjunction between nucleolar stress–induced apoptotic mechanisms and genotoxic stress pathways. This refinement is crucial because it allows the exploitation of the therapeutic vulnerability created by aberrant ribosome biogenesis in cancer cells without the added burden of double strand brakes-induced side effects.

### Ribosome biogenesis impairment by targeting protein-protein interactions

The molecular intricacies of the transitional states involved in the formation of a fully operational ribosome furnish compelling evidence regarding the significance of protein-protein interactions in this biological process. It has been shown that ribosomal proteins released during impaired ribosome biogenesis bind to regulatory proteins such as Mdm2, thereby stabilizing p53. The disruption of the interaction between Mdm2 and tumor-suppressor regulators like RPL5, RPL11, and RPL23 results in p53 activation, providing a clear link between targeting protein‒protein interactions and modulating ribosome biogenesis ^37,38^. Additional research highlights interactions among regulatory proteins, including ARF, NPM, and RPL11 with Mdm2, which govern both ribosome biogenesis and cell cycle control. Concurrently, the disruption of certain protein-protein interfaces may result in the impairment of ribosome biogenesis. The interference of RNA polymerase I interactions with the transcription initiation factor Rrn3, facilitated by the application of peptides, significantly impedes ribosome biogenesis and precipitates apoptosis in neoplastic cells^39^. The perturbation of these protein-protein interactions further substantiates the hypothesis that the modulation of such interfaces can have a profound effect in the ribosome biogenesis pathway ^38^.

We present evidence that ribosome biogenesis can be adversely affected through the precise modulation of protein-protein interactions that take place during the ribosome maturation process. To the best of our understanding, this constitutes the first documented occasion in which ribosome biogenesis has been strategically addressed through the interruption of protein-protein interactions that occur throughout the rRNA maturation pathway, as opposed to perturbing the very initial steps of rRNA synthesis. This revelation broadens the scope of research, given the multitude of protein interactions present within the ribosomal synthesis pathway. The contemporary data concerning the architectures of ribosome maturation intermediates acquired through cryo-electron microscopy presents a pivotal opportunity that facilitates the exploration of novel intervention strategies for the development of agents that disrupt protein-protein interactions.

## METHODS

### Protein production

Plasmid 6N31 (based on pFBOH-MHL) 6N31 was a gift from Cheryl Arrowsmith (Addgene plasmid # 127293 ; http://n2t.net/addgene:127293 ; RRID:Addgene_127293) and encodes human WDR12 β-propeller domain, with a TEV cleavable His6 at the N-terminal domain. Bacmids where obtained using the Bac-to-Bac method by transforming 6N31 in E. coli DH10Bac^™^. LB plates were supplemented with kanamycin (50 μg/ml), tetracycline (12 μg/ml), gentamicin (7 μg/ml), IPTG (40 μg/ml) and X-gal (100 μg/ml). White colonies were selected followed by colony PCR for insert detection. Resulting positive bacmid was transfected into Sf9 insect cells on 6 well plates using supplemented Grace media (Gibco^™^, ref. 11605094) and transfection reaction FuGene® HD (Promega, ref. E2311). For protein production media Sf-900^™^ II SFM (Gibco^™^, ref. 10902088) was used. Protein overespression and purification was made following standard procedures. Typically 300 ml of infected Sf9 were pelleted and sonicated in lysis buffer 50 mM Hepes pH 7,5; 0,5 M NaCl; 10 % (v/v) glycerol; 0,1% (v/v) Triton X-100; 5 mM β-mecaptoethanol. Supernatant was filtered and loaded on a 5ml HisTrap^™^ (Cytiva) column and eluted in a linear gradient of 0-500 mM imidazole using an FPLC system. Fractions were analyzed for purity on a 10% SDS-PAGE.

All peptides were commercially obtained from Synpeptide Co., Ltd., Shanghai, China.

### Interaction assays

Bio-layer interpherometry (BLI) was used to determine the affinity between WDR12 β-propeller domain and the peptides. Sensorgrams were obtained through the immobilization of 50 μg/ml N-terminal biotinylated peptides (Biot-Bop1_439-449_ and Biot-Bop1_405-412_) onto streptavidin biosensors (FortéBio Sartorius) employing analyte concentration ranges spanning from 15 to 50 μM of purified WDR12 β-propeller for the association phase. For each analyte concentration, real time changes in the interference pattern were monitored and plotted overtime. Resulting curves were used to calculate K_D_values using the Blitz Pro 1.2 software.

### Cell culture

The HCT116 human colon cancer line was provided by Dr. Manuel Serrano (AltosLaboratories, Cambridge). The cells grew in a supplemented DMEM with 10% fetal bovine serum, 2 mM of L-glutamine, 150 u/ml of penicillin and 150 μg/ml of streptomycin in an incubator with humidity saturated at 37°C and 5% CO_2_.

### Immunofluorescence of cells in culture

For nucleolin localisation assays, 3 × 10^4^ HCT116 cells per well were seeded in 24-well plates on coverslips pre-treated with laminin at 0.5 µg/ml in PBS for at least 1 hour. The following day, 24h treatments were carried out with 100 µM of the P10hs and TAT peptides and 200 µM of P11hs. Once the treatment was complete, the cells were washed with PBS and fixed with a 4% (w/v) paraformaldehyde solution in PBS for 15 minutes at room temperature. They were then washed with PBS and permeabilised with 1% (w/v) Triton X-100 in PBS for 15 minutes at room temperature. After several washes, the samples were blocked with 0.1% Tween 20 and 4% fetal bovine serum in PBS for 1 hour at room temperature and then incubated with the primary antibody nucleolin C23 (MS-3, sc-8031 Santa Cruz) in blocking solution overnight at 4°C. After washing three times with PBS, the samples were incubated with Alexa Fluor 488 anti-mouse IgG secondary antibody for 1 hour at room temperature. The secondary antibody was washed in the same manner and the nuclei were stained with a PBS solution containing 1µg/ml DAPI (Thermo 62248) for 15 minutes. Finally, after three washes with PBS, the coverslips were mounted on slides using Aqua-Poly/Mount medium (Polysciences). Images were acquired using a Leica SP8 confocal microscope (Leica MicroSystem) and analyzed using ZEN Blue and ImageJ software.

To identify the location of the P10hs and P11hs peptides, HCT116 cells were seeded at 2×10^4^ cells per well in 24-well plates on laminin-treated coverslips. After 24 hours, they were treated with 5 µM FITC-P10hs and FITC-P11hs for 5 hours. They were then fixed for 15 minutes with a 4% (w/v) paraformaldehyde solution in PBS and, after a couple of washes, the nuclei were stained with a solution containing DAPI. They were washed three times with PBS and the coverslips were mounted with Aqua-Poly/Mount mounting medium. Images were acquired using a Leica SP8 confocal microscope (Leica MicroSystem) and analyzed using ZEN Blue and ImageJ software.

### Cell proliferation assays

Cell proliferation was determined using the Cell Proliferation Reagent (WST-1) kit. HCT116 cells were seeded in triplicate in 96-well plates at a density of 2.5 × 10^3^ cells per well. After 24 hours, treatments were performed by adding the different interference peptides at different concentrations: 50, 100, 150 and 200 µM for peptide P10hs, 100, 200, 300, 400, 500 and 600 µM for peptide P11hs, and 200 and 600 µM for peptide TAT. At 24, 48 and 72 hours of treatment with the peptides, 10 µl of WST-1 reagent was added per well and, after incubation for 3 hours at 37°C, the absorbance was measured at 450 nm in the VICTOR2 1420 Multilabel Counter spectrophotometer (Wallac).

### Cell transduction analysis

#### 1. SUnSET assay

HCT116 cells were seeded in 60 mm plates at a density of 5 × 10^5^ per plate, and the following day, treatments were performed with the interfering peptides P10hs and TAT at 100 µM. After 24 hours, 5 µg/ml puromycin was added directly to the culture medium and the cells were incubated for 15 minutes at 37°C. The cells were then washed with PBS, collected, and total protein extracts were prepared. The protein translation rate was assessed by Western blot analysis using the anti-puromycin antibody (clone 12D10, MABE343 from Sigma-Aldrich).

#### 2. Polysomal profile

To perform this assay, HCT116 cells were seeded in 150 mm plates at a density of 7.5 × 10^6^ cells per plate and treated the following day with 100 µM of the P10hs peptide, 200 µM of the P11hs peptide and 200 µM of TAT. Twenty-four hours after treatment and at 80% confluence, 100 µg/ml of cycloheximide (CHX) was added to the cell plates and incubated for 10 minutes at 37°C. After washing with cold PBS + 100 µg/ml CHX, the cells were lifted with trypsin and centrifuged at 500 x g for 5 minutes at 4°C. After removing the supernatant, they were washed with cold PBS + 100 µg/ml CHX, centrifuged again, and the pellet was resuspended in lysis buffer composed of 20 mM Tris-HCl (pH 7.4), 5 mM MgCl_2_, 150 mM NaCl, 1 mM DTT, 100 mg/ml CHX, 40 u/ml RNase inhibitor, protease inhibitors (cOmplete™ EDTA-free Protease Inhibitor Cocktail, Roche), 0.5% Triton X-100 and 0.5% deoxycholate. The extract was centrifuged for 5 min at 2,000 x g at 4°C and the collected supernatant was centrifuged again for 15 min at 14,000 x g at 4°C. Finally, to load the same amount of sample into all sucrose gradients, the OD at 260 nm of the supernatant was measured in a Nanodrop 2000 (Thermo Fisher Scientific), and the equivalent of 400 mg of RNA was diluted to a final volume of 200 µl in lysis buffer.

Next, 10% and 50% sucrose solutions were prepared in a buffer composed of 20 mM Tris-HCl (pH 7.4), 5 mM MgCl_2_, 150 mM NaCl, 1 mM DTT, 100 mg/ml CHX, 40 u/ml RNase inhibitor, and protease inhibitors. The 10%– 50% sucrose linear gradients were prepared in the Gradient Master (Biocomp Instruments) and kept cold until use. The 200 µl samples were carefully deposited into each gradient, and after balancing them with the aid of a balance, they were loaded into the SW40Ti rotor (Beckman Coulter). They were centrifuged in the Beckman Optima LE-80K ultracentrifuge (Beckman Coulter) at 36,000 rpm for 2 hours at 4°C with slow acceleration and deceleration. The gradients were fractionated using a piston gradient fractionator system (Gradient Station, BioComb Model 153) coupled to the Triax (Triax Flow Cell), which allowed the absorbance of the gradient to be measured in real time.

### Caspase activity analysis

HCT116 cells were seeded in 96-well plates at a density of 2 × 10^3^ cells per well and after 24 hours in the incubator, the corresponding peptides were added: 50, 100, 150 and 200 µM of P10hs; 100, 200, 300, 400 and 600 µM of P11hs and 100 and 600 µM of the TAT peptide, with triplicates for each condition tested. After 24 hours of treatment, caspase activity was determined using the Caspase-Glo® 3/7 kit (Promega) following the manufacturer’s instructions. The reagent was added in a 1:1 ratio and after 2 hours at room temperature in the dark, the luminescence of each well was measured in the VICTOR2 1420 Multilabel Counter (Wallac) plate reader, with the luminescence signal being proportional to the caspase activity present in each sample.

### Protein analysis Western Blot

To obtain total protein extracts, cells were collected in PBS and, after centrifuging for 10 minutes at 2,000 x g, the cell pellet was resuspended in RIPA lysis buffer (50 mM Tris-HCl pH 7.5, 150 mM NaCl, 1 mM EGTA, and 1% NP-40) supplemented with protease inhibitors (cOmplete™ EDTA-free Protease Inhibitor Cocktail, Roche) and phosphatase inhibitors (PhosSTOP™ Phosphatase Inhibitor Cocktail Tablets, Roche). After incubating the lysate for 15 minutes on ice, it was centrifuged at 21,000 x g for 10 minutes at 4°C, and the protein concentration in the supernatant was determined using the Bradford colorimetric method (Bio-Rad Protein Assay).

Next, 20 μg of total protein from each sample was separated on 10% SDS-PAGE gels, after denaturing the proteins by incubating them at 95°C for 5 minutes and then transferring them to nitrocellulose membranes (0.45 μm Hybond ECL Nitrocellulose, Amersham) using a semi-dry transfer system. These membranes were blocked with a 5% solution of skimmed milk in 0.1% PBS-Tween for at least 1 hour with gentle agitation and then incubated with the primary antibody overnight at 4°C. After 3 washes with PBS-Tween 0.1%, the membrane was incubated for 1 hour with the corresponding secondary antibody conjugated with peroxidase dissolved in the same blocking solution. To reveal the activity of the HRP enzyme, the chemiluminescent substrate ECL (Pierce ECL Western Blotting Substrate (Thermo Scientific™)) was added, and its luminescence was detected using the ImagenQuant™ LAS 4000 Mini (GE Healthcare) equipment.

The following antibodies were used for the assays: Caspase 3 (sc-271028, Santa Cruz), Caspase 8 (sc-56070, Santa Cruz), Caspase 9 (sc-56073), Phospho-Histone H2A. X (Ser139) (20E3) (#9718, Cell Signaling), Actin (A2066, Sigma), Anti-Rabbit IgG (NA934, Cytiva) and Anti-Mouse IgG (NXA932, Cytiva).

### Flow cytometry

For flow cytometry assays, HCT116 cells were seeded in 6-well plates at a density of 7.5 × 10^4^ cells per well and after 24 hours were treated with 50, 100 and 150 µM P10hs, 200 and 400 µM P11hs, and 150 and 400 µM TAT. As a positive control for apoptosis, one well at the same cell density was treated with 1 µM camptothecin.

The cells were trypsinized after 24 hours of treatment and centrifuged at 300 x g for 10 minutes. The cell pellet was resuspended in 100 µl of binding buffer at a concentration of 1 × 10^6^ cells/ml. Subsequently, 5 µl of Annexin-FITC V and 5 µl of propidium iodide (PI) were added to each tube (FITC Annexin V Apoptosis Detection kit I, BD Biosciences), and after vortexing, they were incubated for 15 minutes at room temperature in the dark, subsequently adding 400 µl of binding buffer to each tube. The fluorescence of 20,000 events was monitored in an LSR Fortessa X-20 flow cytometer (BD Biosciences). Blue (488 nm) and yellow-green (561 nm) lasers were used to measure Annexin-FITC V (λ_ex_ = 488 nm, λ_em_ = 530/515 nm) and PI (λ_ex_ = 561 nm, λ_em_ = 586/515 nm).

## ACKNOWLEDGEMENTS

This research was supported by grant PID2020-120243RB-I00 from the spanish “Ministerio de Ciencia e Innovación”. We thank Tatiana Ramírez Valencia for assistance in BLI assays and Dr Francisco José Iborra for helpful discussion.

